# Roscovitine exacerbates *Mycobacterium abscessus* infection by reducing NADPH oxidase-dependent neutrophil trafficking

**DOI:** 10.1101/2021.07.30.454490

**Authors:** Vincent Le Moigne, Daniela Rodriguez Rincon, Simon Glatigny, Christian M. Dupont, Christelle Langevin, Amel Ait Ali Said, Stephen A. Renshaw, R. Andres Floto, Jean-Louis Herrmann, Audrey Bernut

## Abstract

Persistent neutrophilic inflammation associated with chronic pulmonary infection causes progressive lung injury and eventually death in individuals with cystic fibrosis (CF), a genetic disease caused by bi-allelic mutations in the cystic fibrosis transmembrane conductance regulator (CFTR) gene.

We therefore examined whether Roscovitine, a cyclin-dependent kinase inhibitor that (in other conditions) reduces inflammation while promoting host defence, might provide a beneficial effect in the context of CF.

Herein, using CFTR-depleted zebrafish larvae as an innovative vertebrate model of CF immuno-pathophysiology, combined with murine and human approaches, we sought to determine the effects of Roscovitine on innate immune responses to tissue injury and pathogens in CF condition.

We show that Roscovitine exerts anti-inflammatory and pro-resolution effects in neutrophilic inflammation induced by infection or tail amputation in zebrafish. Roscovitine reduces overactive epithelial ROS-mediated neutrophil trafficking, by reducing DUOX2/NADPH-oxidase activity, and accelerates inflammation resolution by inducing neutrophil apoptosis and reverse migration. Importantly, while Roscovitine efficiently enhances intracellular bacterial killing of *Mycobacterium abscessus* in human CF macrophages *ex vivo*, we found that treatment with Roscovitine results in worse infection in mouse and zebrafish models. By interfering with DUOX2/NADPH oxidase-dependent ROS production, Roscovitine reduces the number of neutrophils at infection sites, and consequently compromises granuloma formation and maintenance, favouring extracellular multiplication of *M. abscessus* and more severe infection.

Our findings bring important new understanding of the immune-targeted action of Roscovitine and have significant therapeutic implications for safety targeting inflammation in CF.

## Introduction

Cystic fibrosis (CF) is a fatal disorder resulting from mutations in the cystic fibrosis transmembrane conductance regulator (CFTR)^1^. The leading causes of premature death in CF individuals is progressive pulmonary injury and respiratory failure caused by mucus obstruction, infections and inflammation^2^.

In CF lungs, impaired CFTR results in airway surface liquid dehydration and collapse of mucociliary clearance, predisposing to recurrent infections with a subsequent hyper-inflammatory profile^2^. CF infections are typified by pathogenic bacteria such as *Pseudomonas aeruginosa, Staphylococcus aureus, Burkholderia cenocepacia* or the non-tuberculous mycobacteria *Mycobacterium abscessus* (Mabs)^3^. In addition, CFTR deficiency results in abnormal activation of macrophage and epithelial cell responses to pathogens^4^, releasing pro-inflammatory mediators, such as IL8 and reactive oxygen species (ROS). This favours the onset of an exuberant influx of neutrophils^4–7^, which nonetheless fails to control infections and worsens lung function^8,9^. Moreover, defects in CFTR impair the ability of neutrophils to undergo apoptosis^10–12^ and reverse migration^7^ leading to increased neutrophil activity and longevity and therefore contribute to sustained pulmonary inflammation^7,12^. Evidence suggests that inflammation may even precede infection in CF aiways^13–15^. Elevated inflammatory markers in the bronchoalveolar lavage fluid of CF infants are found, even in the absence of detectable infection^16^. In particular, we have demonstrated that CFTR dysfunction directly alters the response of epithelial cells to “sterile” injury and leads to exuberant ROS production through the DUOX2/NADPH oxidase, driving an overactive neutrophil response in a CFTR-depleted zebrafish model^7^.

Reducing the deleterious impact of inflammation is therefore an important therapeutic goal in CF^17^. Conventional anti-inflammatory therapies in CF include the use of glucocorticoids or ibuprofen which are potentially effective but associated with significant long term side effects^18^. CFTR modulators have been shown to reduce inflammation, however their high cost and mutation/age restriction preclude widespread use. Antibiotic treatment alone is insufficient to prevent inflammatory lung damage and can induce antimicrobial resistance. Although inflammation is reduced with anti-inflammatory treatment^19^, chronic inflammation remains a consistent feature, indicating a continued need for novel approaches to prevent inflammation-mediated tissue destruction in CF.

One potential and interesting alternative is represented by Roscovitine, an inhibitor of cyclin-dependent kinases (CDK)^20^. In particular, this compound is capable of inducing neutrophil apoptosis^21,22^, accelerating the resolution of inflammation^23–25^. Importantly, Roscovitine has proven beneficial in enhancing apoptosis of neutrophils isolated from CF patients^11^. However, the pro-apoptotic activity of Roscovitine has never been evaluated in *in vivo* models of CF. Roscovitine also exerts anti-inflammatory actions on macrophages^26,27^, eosinophils^28,29^ and lymphocytes^30^. Moreover, Roscovitine enhances bactericidal activity of CF alveolar macrophages^31,32^. However, Roscovitine has not been tested in CF infection models. Roscovitine is currently being evaluated in a phase 2 clinical trial in CF patients infected with *P. aeruginosa*, as a potential anti-pseudomonas therapy https://clinicaltrials.gov/ct2/show/NCT02649751?term=roscovitine&rank=1.

Here, we demonstrate that Roscovitine can restore normal levels of inflammation in a *in vivo* model of CF by *i)* reducing epithelial ROS production-driven neutrophil mobilisation and *ii)* enhancing neutrophil apoptosis and reverse migration. Importantly, beside macrophage-directed bactericidal effect of Roscovitine, we show that Roscovitine promotes an increased susceptibility to Mabs infection *in vivo* by inhibiting DUOX2/NADPH oxidase-dependent neutrophil trafficking. This study represents a clear demonstration of the protective role of DUOX2-mediated ROS production against Mabs infection.

## Methods

Bacterial strains, human cells, mouse and zebrafish lines and detailed methods associated with all procedures below are available in **Supplemental Methodology**.

### Zebrafish experiments

Zebrafish experiments were conducted according to guidelines from the UK Home Office under AWERB and in compliance with the European Union guidelines for handling of laboratory animals.

### Mouse experiments

Mouse procedures were authorised by Ethics Committee A783223 (APAFIS#11465-2016111417574906).

### Macrophage experiments

Primary human macrophages were generated from peripheral blood samples from consented healthy and individuals with CF volunteers (approved by regional ethics approval REC12/WA/0148).

### Quantification and statistical analysis

Statistical analysis was performed using Prism 7.0 (GraphPad Software) and detailed in each Figure legend. ns, not significant (p≥0.05); *p<0.05; **p<0.01; ***p<0.001; ****p<0.0001.

## Results

### Roscovitine rebalances early neutrophil infiltration by epithelial ROS-dependent mechanisms

We first proceeded to examine the potential benefits of Roscovitine in reducing neutrophilic inflammation by exploiting the zebrafish model of sterile inflammation^7,33,34^. In zebrafish larvae, tail fin amputation triggers neutrophil infiltration towards wound, accurately mimicking the kinetics and fates observed in human inflammatory responses^33,35^. In particular, zebrafish neutrophils have the same function as human neutrophils and respond in a similar manner to chemicals, including Roscovitine^23^.

In order to investigate the effect of Roscovitine on neutrophilic response, we exploited the *TgBAC(mpx:EGFP)i114* line harbouring green-fluorescent neutrophils^33^, in normal and CFTR-deficient contexts, using *cftr* morphants (*cftr* MO)^6^ or the knockout *cftr*^sh540^ mutant (*cftr* -/-)^7^. To first address whether Roscovitine influences early neutrophil infiltration, injured-WT and CF larvae were incubated with Roscovitine, or *i)* the NADPH-oxidase blocker Diphenyleneiodonium (DPI), known to inhibit early neutrophil mobilisation^7,36^, *ii)* the pro-resolution drug Tanshinone IIA (TIIA), which does not influence early neutrophil chemotaxis^7,37^ and *iii)* DMSO. Roscovitine treatment, but not TIIA, was able to reduce neutrophil influx in WT and CF injured-fish, effectively rebalancing overactive neutrophil mobilisation in CF to that of WT levels (**Figures 1A-B**). Interestingly, comparative analysis showed similar wound-associated neutrophil number in both DPI- and Roscovitine-treated larvae. Epithelial release of H_2_O_2_, through the DUOX2/NADPH oxidase, is required for the early neutrophil response to injury^7,38,39^. We then investigated the potential anti-oxidative action of Roscovitine on the recruitment of early-arriving neutrophils, by measuring ROS production in injured CF fish. Compared to DMSO-treated animals, microscopy revealed that Roscovitine caused a substantial inhibition of epithelial ROS production, as judged by decreased CellROX fluorescence intensity at the wound (**Figures 1C-D**). This finding suggests that Roscovitine modulates the earliest phase of neutrophil mobilisation to injury in an epithelial oxidase-dependent manner.

**Figure 1.**
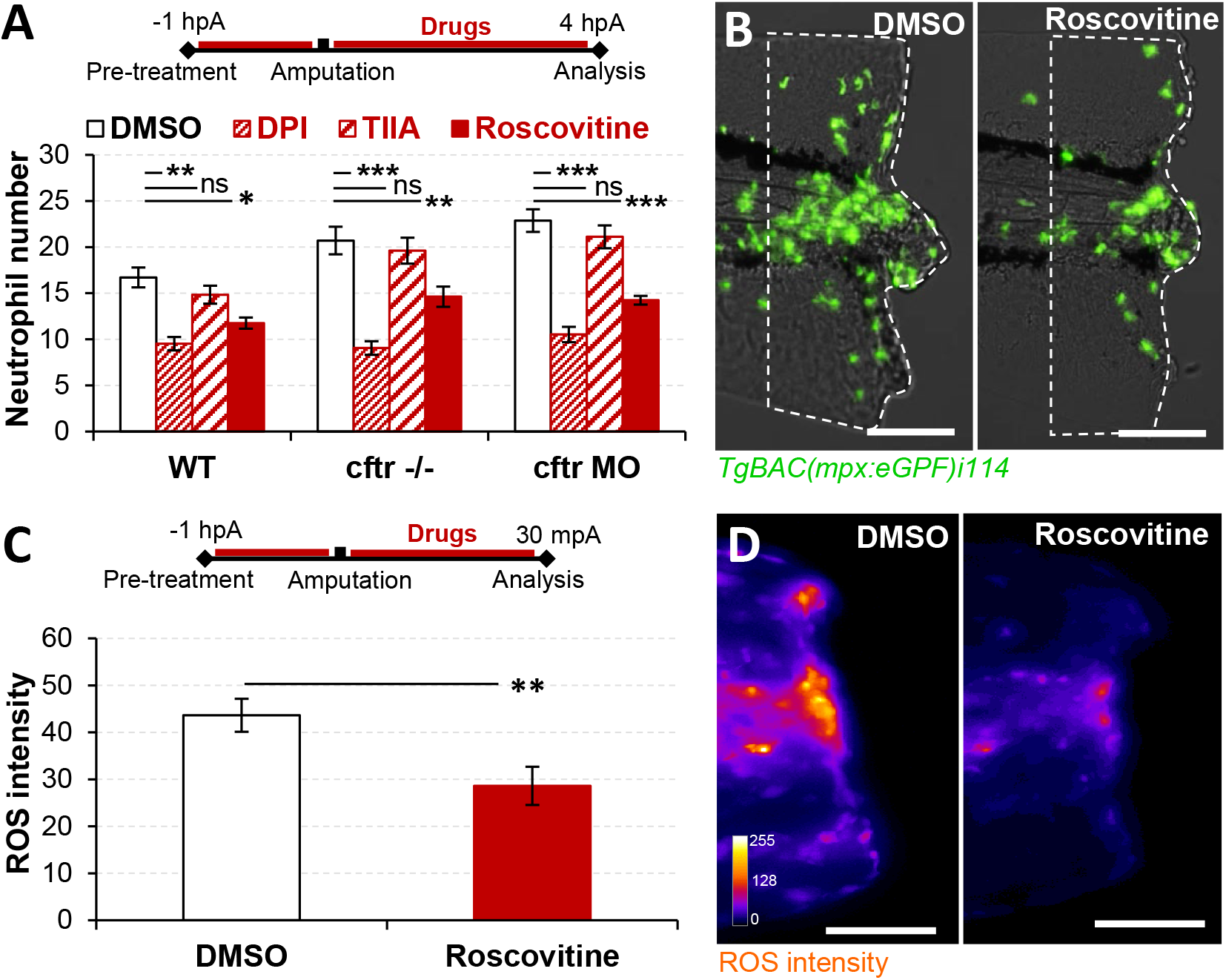
Roscovitine-reduced epithelial oxidative activity rebalances early neutrophil mobilisation at wound in CF zebrafish model. (A-B) WT, *cftr* –/– and *cftr* MO *TgBAC(mpx:EGFP)i114* larvae were pre-treated with of Roscovitine, DPI (as positive control) TIIA (as negative control) or DMSO (as mock control) prior to tail fin amputation procedure, then injured and immediately put back in treatments for 4 h. Neutrophil number at the wound (dotted lines) was observed and enumerated at 4 hpA under a fluorescence microscope. (A) Neutrophil recruitment assay (*n*= 21, Two-Way ANOVA with Dunnett’s post-test, error bars represent SEM). (B) Representative number of neutrophils at wound in Roscovitine-*versus* DMSO-treated *cftr* MO zebrafish (Scale bars, 200 μm). (C-D) *cftr* MO stained with CellROX^®^ to label H_2_O_2_ generation. Means ± SEM ROS intensity (C) and associated pseudocolored photomicrographs (D) of injured tails revealing oxidative activity at 30 min post-amputation (mpA) in *cftr* MO treated with Roscovitine (*n* = 12, Mann Whitney test; Scale bars, 200 μm).

Collectively, these results indicate that Roscovitine reduces CF-associated inflammation by reducing both epithelial oxidase activity and early neutrophil influx to injured tissue in CFTR-depleted zebrafish.

### Roscovitine-driven neutrophil apoptosis and reverse migration accelerate inflammation resolution *in vivo*

CF zebrafish exhibit persistent neutrophilic inflammation after injury^7^. We therefore investigated whether Roscovitine treatment could resolve such a response to initiate regenerative processes.

WT and CF *TgBAC(mpx:EGFP)i114* larvae were injured and, 4 hours later, exposed to Roscovitine or DMSO. Roscovitine reduced established post-wounding neutrophilic inflammation in WT^23^ and CF contexts (**Figures 2A-B**). Pro-resolution events such as local neutrophil apoptosis and migration of neutrophils away from inflamed sites play a critical role to reduce inflammation and restore tissue homeostasis^37,40,41^. We first examined the extent of neutrophil apoptosis *in vivo* in CF zebrafish. Combined confocal imaging and quantification of TUNEL-positive neutrophils showed that CFTR-deficient larvae treated with Roscovitine exhibited enhanced neutrophil apoptosis at wound at 8 hours post-amputation (hpA), compared to their control counterparts (**Figures 2C-D**). Interestingly, Roscovitine induces neutrophil apoptosis more efficiently than TIIA (**Supp 1A**). We then investigated whether Roscovitine could also influence neutrophil retrograde migration by examining and comparing the dynamics of neutrophil reverse migration in DMSO- and Roscovitine-treated larvae using *Tg(mpx:Gal4)sh267;Tg(UAS:Kaede)i222* larvae (**Figure 2E**)^7,42,43^. Remarkably, Roscovitine significantly enhanced neutrophil reverse migration in injured CF fish (**Figures 2F-G**). However, Roscovitine is a much less potent inducer of neutrophil reverse migration than TIIA (**Supp 1B**). Efficient inflammation resolution plays a pivotal role preventing tissue damage, as well as initiating tissue healing and repair^44–46^. The pro-resolution property of Roscovitine, linked to increased neutrophil apoptosis and reverse migration, prompted us to analyse tissue repair potential in zebrafish treated with Roscovitine. Despite evidence of reduced damage to regenerated tissues, our results indicated that defective tissue repair was not reversed by Roscovitine exposure in CF animals (**Supp 2A-B**).

**Figure 2.**
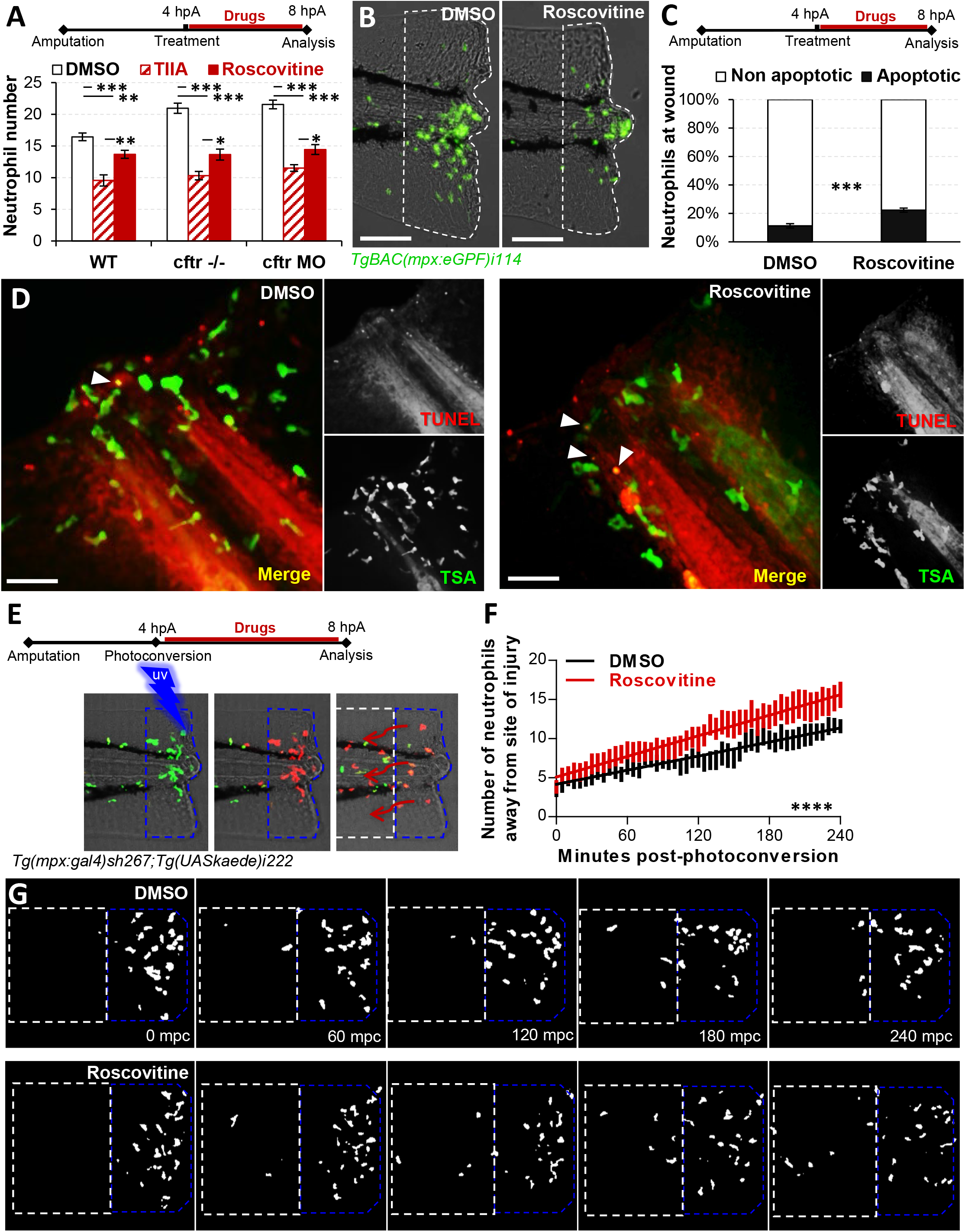
Roscovitine accelerates inflammation resolution *in vivo* both by inducing neutrophil apoptosis and reverse migration. (A-B) Control-Mo or cftr-MO *TgBAC(mpx:EGFP)i114* were injured and treated from 4 hpA with Roscovitine or of TIIA. (A) Neutrophil number at the wound was observed and counted at 8 hpA (n=21, Two-Way ANOVA with Tukey’s multiples comparison test). (B) Representative number of neutrophils remaining at wounds (Scale bars, 200 μm). (C-D) injured-*cftr* MO larvae were treated with Roscovitine from 4 hpA and stained with TUNEL/TSA to label apoptotic cells (C) Neutrophil apoptosis quantification at 8 hpA (n= 15, Fisher *t*-test). (D) Representative confocal pictures of injured tails (Scale bars, 50 μm) revealing the proportion of apoptotic neutrophils at the wound at 8 hpA. (E-F) Reverse-migration in *cftr* MO *Tg(mpx:gal4)sh267;Tg(UASkaede)i222* after Roscovitine treatment. At 4 hpA, neutrophils at site of injury were photoconverted then the numbers of photoconverted cells (red) that migrate away (white dotted box) from the photoconverted area (blue dotted box) were time-lapse imaged and quantified over 4 hours by confocal microscopy (E). (F) Plot showing the number of photoconverted neutrophils leaving the wound over 4 hours post photoconversion (hpc). Line of best fit shown is calculated by linear regression. *P-*value shown is for the difference between the 2 slopes (*n*= 12, performed as 3 independent experiments). (G) Representative confocal imaging of injured tails showing the kinetics of photoconverted neutrophils that move away from the area of injury over inflammation resolution.

Overall, we show that Roscovitine promotes resolution of established neutrophilic inflammation and alleviates inflammatory damage in CFTR-depleted fish by enhancing both neutrophil apoptosis and reverse migration.

### Roscovitine exposure compromises epithelial ROS-dependent neutrophil mobilization during Mabs infection

As neutrophils represent the first line of defence against invading bacteria, including the multi-drug resistant pathogen Mabs^47,48^, we were next interested in determining the effect of Roscovitine on neutrophil responses during Mabs infection, using a zebrafish model of Mabs infection^49,50^. Chemoattraction of neutrophils was assessed by injecting Mabs expressing *tdTomato* into the somite of *TgBAC(mpx:EGFP)i114* larvae as previously described^48^. As shown in **Figures 3A-C**, Roscovitine exposure resulted in a significant reduction in neutrophil mobilisation towards Mabs-infected tissue. Neutrophil chemotaxis is known to require functional epithelial ROS signalling^51^, suggesting this could also account for the Mabs-induced neutrophil response. While injection of Mabs consistently triggers oxidative responses in infected tissues, confocal microscopy showed abnormal oxidative activity in Roscovitine-treated larvae, which causes a substantial inhibition of epithelial ROS generation at the site of infection, as reflected by the decreased CellROX signal (**Figures 3A-B**). Noteworthily, this reduction of ROS production coincides with a reduced number of neutrophils mobilised towards bacilli in fish exposed to Roscovitine (**Figure 3A**). Additionally, confocal examination of Mabs-granuloma, a protective structure improving the control of Mabs infection^48^, revealed an abnormal granuloma architecture in Roscovitine-treated larvae, typified by reduced neutrophil infiltration (**Figure 3C**).

**Figure 3.**
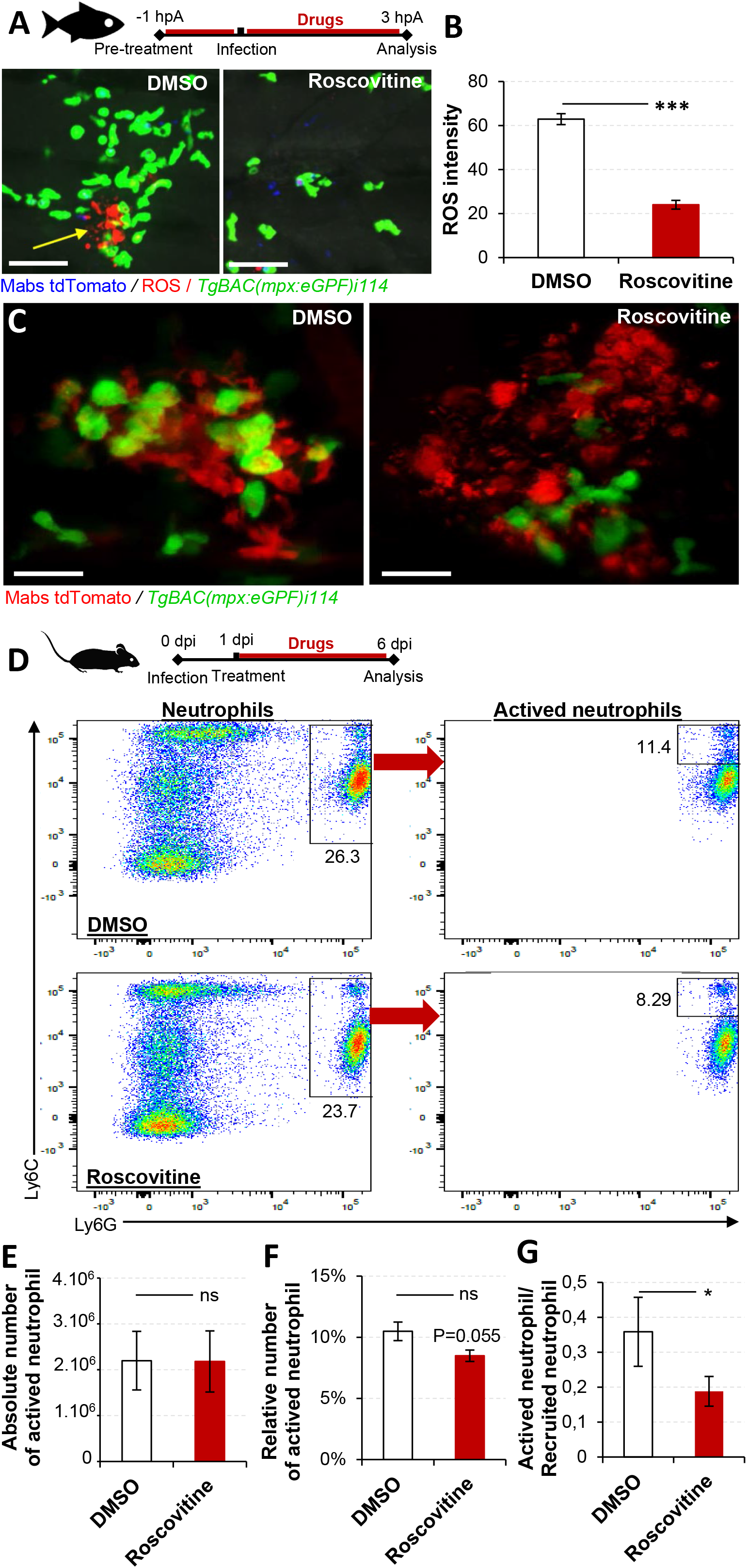
Roscovitine impedes neutrophil trafficking during Mabs infection. (A-B) WT *Tg(mpx:eGFP)i114* larvae were treated with Roscovitine or DMSO then infected into the somite with ≈100 Mabs R expressing dtTomato. Infected larvae are stained with CellROX^®^ to label ROS production. Representative epithelial oxidative response (arrow) and number of neutrophils at infection site in Roscovitine-*versus* DMSO-treated larvae at 3 hours post-infection (hpi) (Scale bars, 75 μm). (B) Means ± SEM ROS intensity at the site of infection (2 hpi, *n* = 8, student t test). (C) Confocal images showing the representative repartition of neutrophil-associated Mabs granuloma in larvae treated with Roscovitine compared with DMSO-exposed animals (Scale bars, 10 μm). (D-G) Mice were intravenously infected with R Mabs then treated with 50 μM Roscovitine or DMSO at 1dpi. At 6 dpi neutrophils are isolated from the lung of mice and analysed by flow cytometry. (D) Representative dot-plots showing the expression of Ly6C^hi^ / Ly6G^hi^ (actived neutrophil) among neutrophils. Graphs showing the mean± SEM absolute (E) and relative (F) number of actived neutrophils, and related ratio of actived neutrophils (G) in lungs (n=5, unpaired Student’s *t* test, representative of 3 independent experiments).

To further support zebrafish experiments, the neutrophil influx and activity were also evaluated in mice infected with Mabs then treated with Roscovitine or DMSO. Neutrophil numbers in lung compartments were enumerated at 6 days post-infection (dpi). As shown in **Figure 3D**, Roscovitine-treated mice exhibited reduced Ly6C^hi^ / Ly6G^hi^ staining, indicating that activated neutrophil amounts has decreased in lung after Roscovitine administration. Reduced relative numbers of activated neutrophil following Roscovitine treatment was confirmed by comparative analysis of cell composition in lung in these mice (**Figures 3E-G**). Of note, no changes in global neutrophil numbers were observed in zebrafish or mice, ensuring that the observed differences did not result from Roscovitine-induced neutropenia (data not shown).

Together these findings indicate that Roscovitine alters neutrophil mobilisation towards Mabs, likely by interfering with epithelial oxidative activity induced by Mabs infection, in addition to the critical role played in granuloma integrity with deleterious consequences such as extracellular mycobacterial multiplication^48^.

### Roscovitine exposure leads to exacerbation of Mabs infection *in vivo*

Neutrophils are dispensable for defence against Mabs infection^48,49,52^. The profound alteration of neutrophil chemotaxis to Mabs caused by Roscovitine, led us to hypothesis that Roscovitine may hamper host defence against Mabs and thus increase susceptibility to Mabs infection.

In order to test whether Roscovitine influences Mabs infection outcomes, intracellular Mabs killing was firstly investigated *ex vivo*, using primary macrophages obtained from both healthy and CF volunteers (**Figure 4A**). Relative luminescent units (RLU) analysis revealed a lower bacterial load in Mabs-infected macrophages treated with Roscovitine compared to vehicle alone at 24 hpi (**Figure 4B**), suggesting that Roscovitine can enhance macrophage Mabs killing in the context of CF. Interestingly, as previously reported, this might depend on the acidification of macrophages^32^, since Roscovitine improves acidification of macrophage lysosomes post Mabs infection, as shown by enhanced lysosomal fusion with intracellular Mabs and increased acidified lysosome numbers in macrophages **(Supp 3A-D)**. To exclude direct Roscovitine-induced Mabs killing as the cause of enhanced mycobacterial clearance in macrophages, we evaluated minimum inhibitory concentrations. None of the Mabs variants showed direct Roscovitine susceptibility (**Table 1**), indicating that this compound has no direct antibacterial activity against Mabs. We demonstrate here that Roscovitine enhances macrophage-mediated intracellular killing of Mabs, likely by improving the lysosomal acidification in macrophages. However, little is known about the effect of Roscovitine on bacterial control *in vivo*.

**Figure 4.**
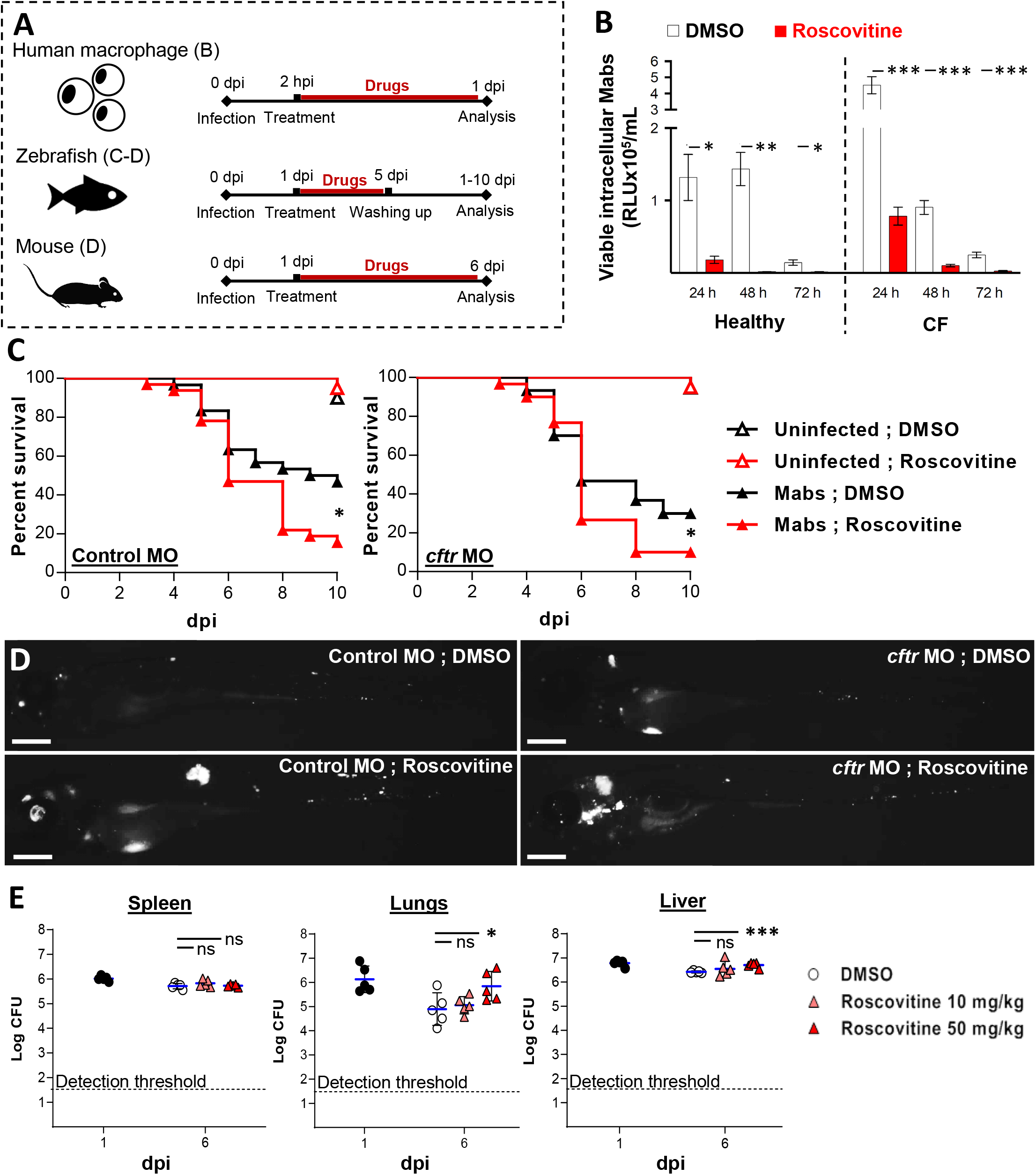
Roscovitine exacerbates Mabs infection in zebrafish and mouse model of infection. (A) The effect of Roscovitine on Mabs infection outcomes was evaluated in primary human macrophage (B), zebrafish (C-D) and mouse (C) model of infection. (B) Monocyte-derived primary human macrophages were infected at a MOI 1:1 with bioluminescent Mabs (Mabs-lux) for 2 hours. Extracellular bacteria were washed off and fresh media containing Roscovitine or DMSO added. At each specified time-point, cells were lysed and viable intracellular bacteria quantified as relative luminescent units (RLU). Roscovitine enhances intracellular Mabs killing in macrophages obtained from both healthy volunteers and CF patients (One-Way ANOVA with Dunnett’s post-test). (C-D) Control MO or *cftr* MO were intravenously infected with ≈100-150 Mabs R expressing *tdTomato*. From one day post-infection (dpi) larvae were treated with Roscovitine or DMSO. (C) Survival analysis of Control MO (left) or cftr MO zebrafish (right). Data are plotted as percentage of surviving animals over a 10 days period (n=30, Mantel-Cox Log-rank test, average of two independent experiments). (D) Representative whole-larvae imaging of Control MO (left) or *cftr* MO zebrafish (right) at 3 dpi (Scale bars, 200 μm). (E) Mice were intravenously infected with Mabs then treated 24 hours later with 10 and 50 μM Roscovitine or DMSO. The surviving bacteria were enumerated after 6 dpi by CFU analysis. Results are expressed as log_10_ units of CFU per organ at 1 (before treatment administration) and 6 dpi (Two-Way ANOVA with Dunnett’s post-test).

Next, to establish whether Roscovitine treatment could affects the control of Mabs infection *in vivo*, zebrafish larvae were intravenously infected with Mabs^50^ (**Figure 4A**). Our results indicated that both control- and *cftr*-MO exposed to Roscovitine displayed hyper-susceptibility to Mabs, correlating with increased larval mortality (**Figure 4C**) and higher bacterial loads (**Figure 4D**). Furthermore, microscopy observations showed that the increase in bacterial loads in Roscovitine-treated fish correlates with replicating extracellular bacteria, translating into increased number of abscesses and cord in the central nervous system of larvae^49^ (**Figure 4D**). This is consistent with a reduced host defence and representative of severe Mabs infection in zebrafish^48^, and thus supports the hypothesis that Roscovitine treatment impedes the control of Mabs. Importantly, a similar impact of Roscovitine upon bacterial load was observed in mice infected with Mabs. Indeed, infected mice treated with Roscovitine displayed reduced ability to clear Mabs (**Figure 4E**) in the first days of infection, likely due to reduced neutrophil activity (**Figures 3D-F**). These phenotypes are in line with the increased bacterial loads in Roscovitine-treated mice infected with *Streptococcus pneumoniae*^53^.

Collectively, these results indicate that despite the favourable impact of Roscovitine on macrophage-mediated killing of Mabs, its activity increases *in vivo* susceptibility to Mabs infection, likely by hampering neutrophil chemotaxis towards infected sites and the nascent granuloma.

### DUOX2/NADPH-oxidase-driven neutrophil recruitment is crucial to control of Mabs *in vivo*

Release of H_2_O_2_ gradients by epithelial cells through DUOX2/NADPH oxidase has been implicated in neutrophil chemotaxis to infected tissues^51^. Our results above suggest that epithelial ROS generation is required for neutrophil mobilization in response to Mabs infection (**Figure 3A**). We therefore investigated whether DUOX2 activity drives neutrophil recruitment to Mabs infection sites. DUOX2/NADPH oxidase was depleted^54^ and the dynamic of neutrophils recruitment examined in *TgBAC(mpx:EGFP)i114* larvae. Inactivation of NADPH oxidase activity though injection of the *duox2* morpholino impaired neutrophil mobilization to the Mabs-infected somite (**Figures 5A-B**). This implies that DUOX2/NADPH oxidase-dependent ROS production is specifically required for early neutrophil chemotaxis towards Mabs. Additionally, confocal imaging underscored reduced number of neutrophil-associated granuloma in the absence of *duox2* signalling (**Figure 5C**). Importantly, loss of DUOX2 correlated with a defective neutrophil trafficking phenotype and abnormal granuloma architecture, similar to the one observed in infected fish treated with Roscovitine (**Figures 3B-C**). To characterise the role of *duox2* in Mabs infection control, both R and S variants were intravenously injected into control- and *duox2-*MO embryos. *duox2* knockdown resulted in a higher susceptibility to Mabs infections, associated with increased larval killing (**Figures 5D-G**) and enhanced bacterial loads, as demonstrated by determination of the fluorescent pixel count (FPC; **Figures 5E-H**) and whole-larvae imaging (**Figures 5F-I**), further substantiating the importance of DUOX2/NADPH oxidase in controlling Mabs infection. Importantly, the increased susceptibility to Mabs infections in absence of DUOX2 activity correlates with enhanced extracellular bacterial multiplication, as evidenced by the higher number of abscesses (**Figures 5J-K**) as well as altered granuloma integrity (**Figures 5L-M**).

**Figure 5.**
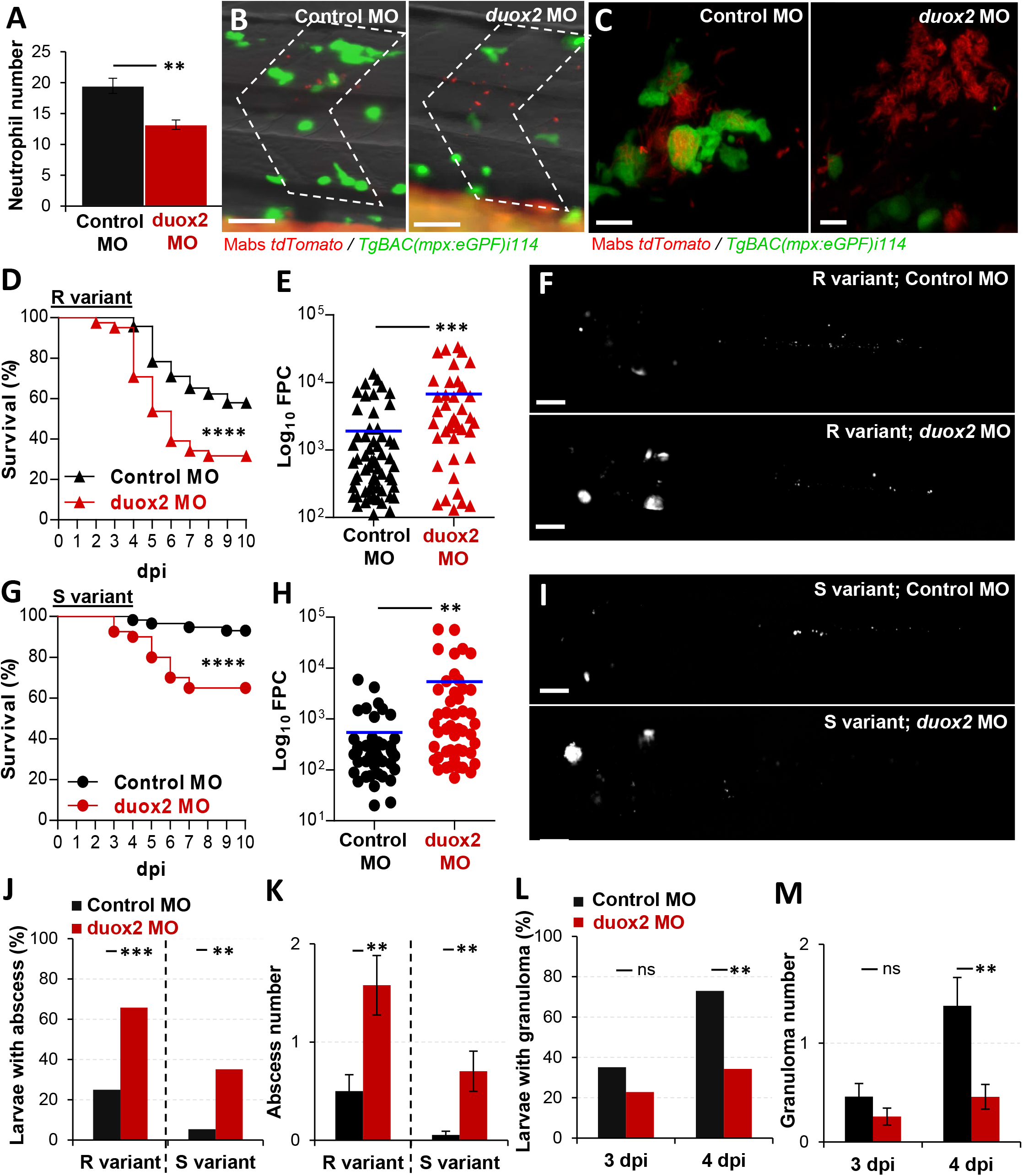
Epithelial oxidative response-dependent recruited neutrophils restricts Mabs infection. (A-B) Control MO or *duox2* MO *Tg(mpx:eGFP)i114* larvae were infected into the somite with ≈100 CFU Mabs S expressing *dtTomato*. (A) Mean ± SEM number of neutrophils mobilized to the infection site at 3 hpi (n= 20, average of two independent experiments) and (B) representative neutrophil-associated site of infection (Scale bars, 75 μm). (C-M) Control MO or *duox2* MO were intravenously infected with ≈100-150 CFU of Mabs R or S expressing *tdTomato*. (C) Confocal images showing the representative repartition of neutrophil-associated *Mabs* granuloma in Control MO *versus duox2* MO (Scale bars, 10 μm). (D and G) Survival analysis of R-(D) or S-infected larvae (G). Data are plotted as percentage of surviving animals over a 10 days period (n=60, Mantel-Cox Log-rank test, Average of three independent experiments). (E and H) Mean fluorescent pixel counts (FPC) of 3 dpi larvae infected by either R (E) or S (H) variants. Results are expressed as log_10_ units of FPC per fish. (F and I) Representative images of R-(F) or S-infected larvae (I) at 3 dpi (Scale bars, 200 μm). (J and K) Percentage of 3 dpi infected larvae with abscess (J) from three independent experiments (n=30) and associated mean ± SEM number of abscess per infected animal (K). (L-M) Kinetic of granuloma formation in whole embryos over a 4-day infection period (L) from three independent experiments (n=30) and associated mean ± SEM number of granuloma per infected animal (M). Statistical significance: Mantel-Cox Log-rank test (D and G), two-tailed unpaired Student’s t test (B, E, H and K), Fisher’s exact test of a contingency table (J and L) or Two-Way ANOVA with Tukey’s multiples comparison test (M).

Together, these results indicate that release of DUOX2/NADPH oxidase-dependent ROS production at the infected sites represents a critical host defence against Mabs and demonstrate that the DUOX2 axis-dependent attraction of neutrophils is instrumental to efficiently contain bacteria within homeostatic granulomas, thereby preventing extracellular mycobacterial spread and limiting subsequent acute infection and larval mortality.

## Discussion

Overactive neutrophil activity has been directly correlated with the onset of bronchiectasis and airway damage in CF, which in term causes lung function impairment and eventually death of people with CF. Thus, reducing the impact of neutrophil inflammation-mediated lung damage is a major concern in CF.

Among the attractive and innovative molecules to target pathways that are specific of the CF lung pathophysiology, Roscovitine shows multiple beneficial proprieties. In particular, Roscovitine stimulates macrophage bactericidal activity^32^ and promotes neutrophil apoptosis^11^ *ex vivo* in models of CF, suggesting that Roscovitine might simultaneously enhance bacterial killing and promote inflammation resolution, therefore prevent subsequent infectious and inflammatory lung damage in CF. However, evaluating Roscovitine in a CF animal model of infection or inflammation was awaited.

Here, moving from *ex vivo* through *in vivo* models of infection or inflammation, in both normal and CF conditions, we sought to determine the effect of Roscovitine on neutrophilic inflammation and how its activity influences the outcomes of infection and inflammation. Our findings indicate that Roscovitine exerts anti-inflammatory and pro-resolution effects in neutrophil response elicited by either Mabs infection or sterile injury. The proposed mechanism by which Roscovitine influences neutrophil trafficking suggests a reduced epithelial ROS burden due to its inhibiting property on DUOX2/NADPH-oxidase.

Whereas previous studies did not investigate Roscovitine effects early after induction of inflammation, our results reveal that Roscovitine especially attenuated neutrophil mobilisation rapidly after infection or injury. Importantly, our findings show that diminished neutrophil response coincided with a reduced epithelial oxidative activity in CF zebrafish treated with Roscovitine. This concurs with described reduced ROS production after Roscovitine treatment in a carrageenan-induced pleurisyin mouse model of inflammation^55^. Several mechanisms could be proposed to explain the action of Roscovitine on epithelial oxidative response, including a down-regulation of calcium release^56^, NF-κB^26^ or TNFα^55^ expression, as well as direct inhibition of DUOX2/NADPH-oxidase. Neutrophil mobilisation being predominantly elicited by DUOX2-mediated epithelial H_2_O_2_^36,57^, our data suggest that by rebalancing epithelial ROS production, Roscovitine could be able to regulate early neutrophil mobilisation towards infected or inflamed tissue in CF.

Neutrophil apoptosis is impaired in CF^7,58^ and can be reversed by Roscovitine in CF patient-derived neutrophils^11^. Furthermore, here we show that Roscovitine is able to induce *in vivo* apoptosis in CF zebrafish neutrophils. This study represents the first demonstration of the pro-apoptotic action of Roscovitine on neutrophils in an *in vivo* model of inflammation in the context of CFTR deficiency. CF-related inflammation is also determined by alterations in neutrophil reverse migration *in vivo*^7^. Reverse migration of neutrophil plays a crucial role in the resolution of inflammation in CF, since restoring this process using TIIA significantly rebalance neutrophil response in CFTR-depleted zebrafish^7^. Here we show for the first time, that Roscovitine can acts on CF zebrafish to restore the reverse migration ability of neutrophils, uncovering a new potential therapeutic mechanism for Roscovitine to drive inflammation resolution in CF. The mechanisms by which Roscovitine influences neutrophil reverse migration is particularly intriguing and deserve further attention.

CF zebrafish show impaired tissue regeneration after tail-fin amputation^7^, in part due to an unresolved neutrophilic inflammation, and which can be restored by pharmacological manipulation of neutrophil responses using TIIA^7^. Interestingly, while Roscovitine profoundly alleviates neutrophilic inflammation, our experiments show that Roscovitine does not improve tissue repair in injured fish. Possible explanations for this finding include the following: (i) Roscovitine inhibits proteins CDK^20^ and p38MAKP^59^, as well as epithelial ROS production : all these pathways are pivotal in the activation of regenerative processes; (ii) blocking CDK9 using Roscovitine delayed macrophage recruitment to injury^60^, an important cell population in the processes of tissue repair^61^; (iii) in contrast to TIIA, Roscovitine preferentially directs the neutrophil towards apoptosis rather than reverse migration. Following Roscovitine treatment, the large amount of apoptotic neutrophils generated could interfere with the efferocytosis potential of macrophages and thus might exert a prolonged local pro-inflammatory state delaying tissue repair.

With the slow development of new treatments and since Roscovitine is readily available and well-tolerated^62^, these findings could have significant therapeutic implications for potently targeting inflammation in CF lung disease, and thus may support currently therapeutic strategies or could be an alternative to existing anti-inflammatory approaches. These data also suggest Roscovitine might have beneficial effects on the pancreas destruction and CF-related diabetes^63^ or gastrointestinal and colorectal cancers in CF^64,65^. While CF is principally characterised by pulmonary infection and inflammation, intestinal disruption involving chronic inflammation is also a frequent feature^64^. In CF, epithelial surfaces produce an increased ROS burden^7^ with potential genotoxic consequences. While ROS are directly mutagenic to DNA, H_2_O_2_ produced in epithelia is a potent chemoattractant source for neutrophils, driving local inflammation^36^, itself a known driver of tumourigenesis^66^. Moreover, ROS production is also a proliferative signal in many epithelial cell types^67^. Interestingly, Duox2 knockout significantly alleviate intestinal inflammation in a mouse model of ileocolitis^68^, suggesting that targeting DUOX2-mediated ROS production might show promise in the treatment of gastrointestinal cancer in people with CF. Firstly known for its anti-cancer properties, Roscovitine is currently being tested in several phase I and II clinical trials against human cancers^69^. So, by restoring normal level of inflammation in CF, Roscovitine might also, by reducing cell proliferation, epithelial ROS-mediated mutagenesis and inflammation, prevent cancer in CF patients.

Mabs infections are associated with severe pneumonia and accelerated inflammatory lung damage in CF patients^70,71^. In line with results previously obtained^31,32^, Roscovitine reduces intracellular bacterial loads in both WT and CF macrophages infected with Mabs, likely by enhancing their ability to kill bacteria. As intracellular bacterial destruction by professional phagocytes is crucial to control Mabs infection^6,72^, perhaps stimulating antibacterial activity using Roscovitine and thereby precluding the establishment of an acute infection could be a therapeutic strategy in CF-related Mabs infection. Roscovitine stimulates macrophage bactericidal activity by restoring intra-phagolysosome acidic pH^31,32^ (which is abnormally high in CF macrophages^73^). Having shown that professional phagocytes acidify phagosomes to efficiently control Mabs^72,74,75^, Roscovitine-mediated intra-phagosomal acidification could account for the Mabs infection phenotype. Interestingly, Roscovitine was found to inhibit Nox2-mediated ROS production in nociceptive neurons through the blockade of Cdk5^55^. Nox2-mediated ROS production in macrophages and neutrophils is another important antibacterial actor against Mabs^6^. These results could suggest that phagosomal acidification is a more potent microbicidal mechanism against Mabs than ROS activity in phagocytes. At this stage, the differential importance of acidic and oxidative defences in the control of Mabs remains to be firmly established. It will be interesting to see whether Roscovitine influences oxidative responses against Mabs. Answering these questions will provide evidence on the most interesting antibacterial mechanisms that could be enhanced therapeutically to better deal with Mabs infections. In addition, whether Roscovitine influences the antibacterial defence of neutrophils has not yet been tested and remains to be addressed.

Unexpectedly, while Roscovitine was able to enhance Mabs killing *ex vivo*, a substantial exacerbation of Mabs infection was found in mice and zebrafish treated with Roscovitine. In particular, Mabs-infected zebrafish rapidly succumbed when exposed to Roscovitine in both WT and CF conditions. Hyper-susceptibility to Mabs due to the Roscovitine exposure is associated with increased extracellular Mabs multiplication and abnormal granuloma maintenance which are representative of a profound impairment in Mabs control^48,49^. Importantly, this increased susceptibility to Mabs coincides with reduced neutrophil mobilization and activity towards infected compartments in mouse and zebrafish. Our previous work highlighted the critical role of neutrophils in the control of Mabs infection by phagocytosing and killing bacilli^47,76^ and by favouring the formation of granulomas able to restrict extracellular multiplication of Mabs^77^. Zebrafish failed to mount a normal epithelial oxidative response to pathogens when treated with Roscovitine, strongly suggesting that Roscovitine affects ROS-driven chemotaxis guiding neutrophils to the nascent granulomas, potentially promoting extracellular Mabs growth and thereby an acute infection.

Although studies postulated that infection-associated neutrophil recruitment is dispensable to epithelial ROS production^78^, we demonstrate the capacity of neutrophils to migrate in DUOX2-derived ROS dependant manner in response to Mabs, that would be directly involved in the formation of protective granulomas. This result shows for the first time that host-derived epithelial ROS signalling, mediated by DUOX2/NADPH oxidase, can prime neutrophil chemotaxis to Mabs infection and therefore defines a critical role for DUOX2 activity in the control of Mabs infection. As a consequence, oxidative activity blockade by Roscovitine increases the risk of impeding host innate immune response and therefore promote an overwhelming Mabs infection. However, since Roscovitine showed enhanced efficacy in combination with other existing therapeutics such as CFTR modulators^31^, Roscovitine will likely diminish the severity of inflammatory lung injury driven by microbial components, host inflammatory mediators as well as genetic defect in CFTR, and accelerated recovery in the context of antibiotic therapy in CF patients.

In addition, while apoptosis is essential for neutrophil shutdown and initiating inflammation resolution, the reduced number of neutrophils due to the pro-apoptotic Roscovitine action may also affects the ability of immune system to efficiently respond to Mabs infection. In contrast, reverse-migrated neutrophils were found able to mount a response to *S. aureus* infection *in vivo*^79^. At this stage, the role of neutrophil reverse migration in the process of infection and inflammation in CF remains to be fully characterised. Reverse migration could have the potential to be deleterious, allowing localised infection or inflammation to disseminate^80^. Alternatively, encouraging neutrophil egress from infected or inflamed sites could serve as a pro-resolving mechanism^7,37^. Answering these questions in CF pulmonary disease will determine how best to harness apoptosis or reverse migration for therapeutic purposes to drive inflammation resolution while minimizing the risk of impaired innate immunity in people with CF.

To conclude, CFTR mutations affect mucus properties, inflammatory processes and antibacterial defences. These different aspects are intertwined: treating one of these features has consequences on the other two. Given its anti-oxidative action, the application of Roscovitine in CF could induce counterproductive and needs therefore to be further studied.

## Acknowledgments

This study was supported by the European Community’s Horizon 2020-Research and Innovation Framework Program (H2020-MSCA-IF-2016) under the Marie-Curie IF CFZEBRA (751977) and by the FRM -Espoirs de la Recherche-Program (ARF201909009156) to AB, the UK CF Trust Strategic Research Centre award (SRC 002) and the Wellcome Trust 107032AIA to RAF and DR-R, an MRC Programme Grant (MR/M004864/1) to SAR, UK CF Trust workshop funding (160161) and UK CF Trust Strategic Research Centre award (SRC018) to AB, SAR and RAF. This work was supported by the AMR cross-council funding from the MRC to the SHIELD consortium “Optimising Innate Host Defence to Combat Antimicrobial Resistance” (MRNO2995X/1) to AB and SAR.

We would like to thank Laurent Meijer (ManRos Therapeutics, Roscoff, France) for kindly providing us with Roscovitine. We acknowledge the Bateson Centre aquarium staff (University of Sheffield), the IERP-Inrea aquarium staff (Paris-Saclay University) and the CRBM-CNRS and LPHI-CNRS fish facilities (University of Montpellier) for zebrafish maintenance and care. We thank the IERP-Inrea (Paris-Saclay University) and Laurent Kremer (IRIM-CNRS, Montpellier) for giving access to their injection platforms. We thank the 2CARE mouse facility at the University of Versailles Saint-Quentin. We wish to thank the CYMAGES and the Wolfson Light Microscopy ((MRC grant (G0700091) and Wellcome Trust grant (GR077544AIA)) facilities. We also thank the University of Sheffield, the University of Cambridge, the University of Montpellier and the University of Versailles Saint-Quentin for support. Finally, we would like to thank Aurore Desquesnes, Catherine Loynes, David Drew and Chira Rizzo for technical assistance.

## Authorship Contributions

AB conceived the study and wrote the manuscript with input from SAR, RAF and J-LH. RAF, J-LH and AB designed experiments and analysed data. RAF, J-LH and AB guided and supervised the work. CL provided zebrafish tools. VLM, DR-R, SG, C-MD, AAAS and AB performed experiments. All authors contributed to the article and approved the submitted version.

## References

1. Gadsby DC, Vergani P, Csanády L. The ABC protein turned chloride channel whose failure causes cystic fibrosis. Nature. 2006;440(7083):477–483.

2. Elborn JS. Cystic fibrosis. Lancet. 2016;388(10059):2519–2531.

3. Lyczak JB, Cannon CL, Pier GB. Lung infections associated with cystic fibrosis. Clin. Microbiol. Rev. 2002;15(2):194–222.

4. Döring G, Gulbins E. Cystic fibrosis and innate immunity: How chloride channel mutations provoke lung disease. Cell. Microbiol. 2009;11(2):208–216.

5. Cantin AM, Hartl D, Konstan MW, Chmiel JF. Inflammation in cystic fibrosis lung disease: Pathogenesis and therapy. J. Cyst. Fibros. 2015;14(4):419–430.

6. Bernut A, Dupont C, Ogryzko N V., et al. CFTR Protects against Mycobacterium abscessus Infection by Fine-Tuning Host Oxidative Defenses. Cell Rep. 2019;26(7):1828-1840.e4.

7. Bernut A, Loynes CA, Floto RA, Renshaw SA. Deletion of cftr Leads to an Excessive Neutrophilic Response and Defective Tissue Repair in a Zebrafish Model of Sterile Inflammation. Front. Immunol. 2020;11:.

8. Downey DG, Bell SC, Elborn JS. Neutrophils in cystic fibrosis. Thorax. 2009;64(1):81–88.

9. McCarthy C, Emmet O’Brien M, Pohl K, Reeves EP, McElvaney NG. Airway inflammation in cystic fibrosis. Eur. Respir. Monogr. 2014;64:14–31.

10. Dibbert B, Weber M, Nikolaizik WH, et al. Cytokine-mediated Bax deficiency and consequent delayed neutrophil apoptosis: A general mechanism to accumulate effector cells in inflammation. Proc. Natl. Acad. Sci. U. S. A. 1999;96(23):.

11. Moriceau S, Lenoir G, Witko-Sarsat V. In cystic fibrosis homozygotes and heterozygotes, neutrophil apoptosis is delayed and modulated by diamide or roscovitine: Evidence for an innate neutrophil disturbance. J. Innate Immun. 2010;2(3):260–266.

12. Gray RD, Hardisty G, Regan KH, et al. Delayed neutrophil apoptosis enhances NET formation in cystic fibrosis. Thorax. 2018;73(2):134–144.

13. Khan TZ, Wagener JS, Bost T, et al. Early pulmonary inflammation in infants with cystic fibrosis. Am. J. Respir. Crit. Care Med. 1995;151(4):1075–1082.

14. Armstrong DS, Grimwood K, Carlin JB, et al. Lower airway inflammation in infants and young children with cystic fibrosis. Am. J. Respir. Crit. Care Med. 1997;

15. Corvol H, Fitting C, Chadelat K, et al. Distinct cytokine production by lung and blood neutrophils from children with cystic fibrosis. Am. J. Physiol. - Lung Cell. Mol. Physiol. 2003;284(6 28-6):.

16. Verhaeghe C, Delbecque K, de Leval L, Oury C, Bours V. Early inflammation in the airways of a cystic fibrosis foetus. J. Cyst. Fibros. 2007;6(4):304–308.

17. Mitri C, Xu Z, Bardin P, et al. Novel Anti-Inflammatory Approaches for Cystic Fibrosis Lung Disease: Identification of Molecular Targets and Design of Innovative Therapies. Front. Pharmacol. 2020;11:.

18. Lands LC, Stanojevic S. Oral non-steroidal anti-inflammatory drug therapy for lung disease in cystic fibrosis. Cochrane Database Syst. Rev. 2019;2019(9):.

19. Hisert KB, Heltshe SL, Pope C, et al. Restoring cystic fibrosis transmembrane conductance regulator function reduces airway bacteria and inflammation in people with cystic fibrosis and chronic lung infections. Am. J. Respir. Crit. Care Med. 2017;195(12):1617–1628.

20. Meijer L, Borgne A, Mulner O, et al. Biochemical and cellular effects of roscovitine, a potent and selective inhibitor of the cyclin-dependent kinases cdc2, cdk2 and cdk5. Eur. J. Biochem. 1997;243(1–2):527–536.

21. Rossi AG, Sawatzky DA, Walker A, et al. Cyclin-dependent kinase inhibitors enhance the resolution of inflammation by promoting inflammatory cell apoptosis. Nat. Med. 2006;12(9):1056–1064.

22. Leitch AE, Riley NA, Sheldrake TA, et al. The cyclin-dependent kinase inhibitor R-roscovitine down- regulates Mcl-1 to override pro-inflammatory signalling and drive neutrophil apoptosis. Eur. J. Immunol. 2010;40(4):.

23. Robertson JD, Ward JR, Avila-Olias M, Battaglia G, Renshaw SA. Targeting Neutrophilic Inflammation Using Polymersome-Mediated Cellular Delivery. J. Immunol. 2017;198(9):3596–3604.

24. Koedel U, Frankenberg T, Kirschnek S, et al. Apoptosis is essential for neutrophil functional shutdown and determines tissue damage in experimental pneumococcal meningitis. PLoS Pathog. 2009;5(5):.

25. Cartwright JA, Lucas CD, Rossi AG. Inflammation resolution and the induction of granulocyte apoptosis by cyclin-dependent kinase inhibitor drugs. Front. Pharmacol. 2019;10(FEB):

26. Jhou RS, Sun KH, Sun GH, et al. Inhibition of cyclin-dependent kinases by olomoucine and roscovitine reduces lipopolysaccharide-induced inflammatory responses via down-regulation of nuclear factor κb. Cell Prolif. 2009;42(2):141–149.

27. Du J, Wei N, Guan T, et al. Inhibition of CDKS by roscovitine suppressed LPS-induced • NO production through inhibiting NFκB activation and BH4 biosynthesis in macrophages. Am. J. Physiol. - Cell Physiol. 2009;297(3):.

28. Farahi N, Uller L, Juss JK, et al. Effects of the cyclin-dependent kinase inhibitor R-roscovitine on eosinophil survival and clearance. Clin. Exp. Allergy. 2011;41(5):673–687.

29. Duffin R, Leitch AE, Sheldrake TA, et al. The CDK inhibitor, R-roscovitine, promotes eosinophil apoptosis by down-regulation of Mcl-1. FEBS Lett. 2009;583(15):.

30. Yoshida H, Kotani H, Kondo T, et al. CDK inhibitors suppress Th17 and promote iTreg differentiation, and ameliorate experimental autoimmune encephalomyelitis in mice. Biochem. Biophys. Res. Commun. 2013;435(3):.

31. Shrestha CL, Zhang S, Wisniewski B, et al. (R)-Roscovitine and CFTR modulators enhance killing of multi-drug resistant Burkholderia cenocepacia by cystic fibrosis macrophages. Sci. Rep. 2020;10(1):.

32. Riazanski V, Gabdoulkhakova AG, Boynton LS, et al. TRPC6 channel translocation into phagosomal membrane augments phagosomal function. Proc. Natl. Acad. Sci. U. S. A. 2015;112(47):E6486– E6495.

33. Renshaw SA, Loynes CA, Trushell DMI, et al. Atransgenic zebrafish model of neutrophilic inflammation. Blood. 2006;108(13):3976–3978.

34. Renshaw SA, Loynes CA, Elworthy S, Ingham PW, Whyte MKB. Modeling inflammation in the Zebrafish: How a fish can help us understand lung disease. Exp. Lung Res. 2007;33(10):549–554.

35. Henry KM, Loynes CA, Whyte MKB, Renshaw SA. Zebrafish as a model for the study of neutrophil biology. J. Leukoc. Biol. 2013;94(4):633–642.

36. Niethammer P, Grabher C, Look AT, Mitchison TJ. A tissue-scale gradient of hydrogen peroxide mediates rapid wound detection in zebrafish. Nature. 2009;459(7249):996–9.

37. Robertson AL, Holmes GR, Bojarczuk AN, et al. A zebrafish compound screen reveals modulation of neutrophil reverse migration as an anti-inflammatory mechanism. Sci. Transl. Med. 2014;6(225):.

38. Niethammer P, Grabher C, Look AT, Mitchison TJ. A tissue-scale gradient of hydrogen peroxide mediates rapid wound detection in zebrafish. Nature. 2009;459(7249):996–999.

39. Gamaley IA, Klyubin I V. Roles of reactive oxygen species: Signaling and regulation of cellular functions. Int. Rev. Cytol. 1999;188:203–255.

40. Buckley CD, Gilroy DW, Serhan CN, Stockinger B, Tak PP. The resolution of inflammation. Nat. Rev. Immunol. 2013;13(1):59–66.

41. De Oliveira S, Rosowski EE, Huttenlocher A. Neutrophil migration in infection and wound repair: Going forward in reverse. Nat. Rev. Immunol. 2016;16(6):378–391.

42. Elks PM, Van Eeden FJ, Dixon G, et al. Activation of hypoxia-inducible factor-1α (hif-1α) delays inflammation resolution by reducing neutrophil apoptosis and reverse migration in a zebrafish inflammation model. Blood. 2011;118(3):712–722.

43. Holmes GR, Anderson SR, Dixon G, et al. Repelled from the wound, or randomly dispersed? Reverse migration behaviour of neutrophils characterized by dynamic modelling. J. R. Soc. Interface. 2012;9(77):3229–3239.

44. Karin M, Clevers H. Reparative inflammation takes charge of tissue regeneration. Nature. 2016;529(7586):307–315.

45. Mescher AL, Neff AW, King MW. Inflammation and immunity in organ regeneration. Dev. Comp. Immunol. 2017;66:98–110.

46. Eming SA, Wynn TA, Martin P. Inflammation and metabolism in tissue repair and regeneration. Science (80-.). 2017;356(6342):1026–1030.

47. Malcolm KC, Caceres SM, Pohl K, et al. Neutrophil killing of Mycobacterium abscessus by intra- and extracellular mechanisms. PLoS One. 2018;13(4):.

48. Bernut A, Nguyen-Chi M, Halloum I, et al. Mycobacterium abscessus-Induced Granuloma Formation Is Strictly Dependent on TNF Signaling and Neutrophil Trafficking. PLoS Pathog. 2016;12(11):.

49. Bernut A, Herrmann JL, Kissa K, et al. Mycobacterium abscessus cording prevents phagocytosis and promotes abscess formation. Proc. Natl. Acad. Sci. U. S. A. 2014;111(10):.

50. Bernut A, Dupont C, Sahuquet A, et al. Deciphering and imaging pathogenesis and cording of Mycobacterium abscessus in zebrafish embryos. J. Vis. Exp. 2015;2015(103):.

51. Brothers KM, Gratacap RL, Barker SE, et al. NADPH Oxidase-Driven Phagocyte Recruitment Controls Candida albicans Filamentous Growth and Prevents Mortality. PLoS Pathog. 2013;9(10):.

52. Malcolm KC, Caceres SM, Pohl K, et al. Neutrophil killing of Mycobacterium abscessus by intra- and extracellular mechanisms. PLoS One. 2018;13(4):.

53. Hoogendijk AJ, Roelofs JJTH, Duitman JW, et al. R-roscovitine reduces lung inflammation induced by Lipoteichoic acid and Streptococcus pneumoniae. Mol. Med. 2012;

54. Huang C, Niethammer P. Tissue Damage Signaling Is a Prerequisite for Protective Neutrophil Recruitment to Microbial Infection in Zebrafish. Immunity. 2018;48(5):1006-1013.e6.

55. Sandoval R, Lazcano P, Ferrari F, et al. TNF-α increases production of reactive oxygen species through Cdk5 activation in nociceptive neurons. Front. Physiol. 2018;9(FEB):

56. Tamma G, Ranieri M, Di Mise A, et al. Effect of roscovitine on intracellular calcium dynamics: Differential enantioselective responses. Mol. Pharm. 2013;10(12):.

57. Galli F, Battistoni A, Gambari R, et al. Oxidative stress and antioxidant therapy in cystic fibrosis. Biochim. Biophys. Acta - Mol. Basis Dis. 2012;1822(5):690–713.

58. McKeon DJ, Condliffe AM, Cowburn AS, et al. Prolonged survival of neutrophils from patients with ΔF508 CFTR mutations. Thorax. 2008;63(7):660–661.

59. Kim J-E, Park H, Choi S-H, Kong M-J, Kang T-C. Roscovitine Attenuates Microglia Activation and Monocyte Infiltration via p38 MAPK Inhibition in the Rat Frontoparietal Cortex Following Status Epilepticus. Cells. 2019;8(7):746.

60. Hoodless LJ, Lucas CD, Duffin R, et al. Genetic and pharmacological inhibition of CDK9 drives neutrophil apoptosis to resolve inflammation in zebrafish in vivo. Sci. Rep. 2016;5:.

61. Nguyen-Chi M, Laplace-Builhé B, Travnickova J, et al. TNF signaling and macrophages govern fin regeneration in zebrafish larvae. Cell Death Dis. 2017;8(8):e2979.

62. Leven C, Schutz S, Audrezet MP, et al. Non-linear pharmacokinetics of oral roscovitine (Seliciclib) in cystic fibrosis patients chronically infected with pseudomonas aeruginosa: A study on population pharmacokinetics with monte carlo simulations. Pharmaceutics. 2020;12(11):.

63. J E. Symposium Summaries. Pediatr. Pulmonol. 2018;53(S2):S36–S147.

64. Abraham JM, Taylor CJ. Cystic Fibrosis & disorders of the large intestine: DIOS, constipation, and colorectal cancer. J. Cyst. Fibros. 2017;16:S40–S49.

65. Starr TK, Allaei R, Silverstein KAT, et al. A transposon-based genetic screen in mice identifies genes altered in colorectal cancer. Science (80-.). 2009;

66. Hanahan D, Weinberg RA. Hallmarks of cancer: The next generation. Cell. 2011;144(5):646–674.

67. Pinto MCX, Kihara AH, Goulart VAM, et al. Calcium signaling and cell proliferation. Cell. Signal. 2015;27(11):2139–2149.

68. Chu FF, Esworthy RS, Doroshow JH, et al. Deficiency in Duox2 activity alleviates ileitis in GPx1- and GPx2-knockout mice without affecting apoptosis incidence in the crypt epithelium. Redox Biol. 2017;11:.

69. Cicenas J, Kalyan K, Sorokinas A, et al. Roscovitine in cancer and other diseases. Ann. Transl. Med.2015;3(10):.

70. Catherinot E, Roux AL, Macheras E, et al. Acute respiratory failure involving an R variant of mycobacterium abscessus. J. Clin. Microbiol. 2009;47(1):271–274.

71. Esther CR, Esserman DA, Gilligan P, Kerr A, Noone PG. Chronic Mycobacterium abscessus infection and lung function decline in cystic fibrosis. J. Cyst. Fibros. 2010;9(2):117–123.

72. Roux AL, Viljoen A, Bah A, et al. The distinct fate of smooth and rough Mycobacterium abscessus variants inside macrophages. Open Biol. 2016;6(11):.

73. Di A, Brown ME, Deriy L V., et al. CFTR regulates phagosome acidification in macrophages and alters bactericidal activity. Nat. Cell Biol. 2006;8(9):933–944.

74. Laencina L, Dubois V, Le Moigne V, et al. Identification of genes required for Mycobacterium abscessus growth in vivo with a prominent role of the ESX-4 locus. Proc. Natl. Acad. Sci. U. S. A. 2018;115(5):E1002–E1011.

75. Dubois V, Viljoen A, Laencina L, et al. MmpL8MAB controls Mycobacterium abscessus virulence and production of a previously unknown glycolipid family. Proc. Natl. Acad. Sci. U. S. A. 2018;115(43):E10147–E10156.

76. Bernut A, Le Moigne V, Lesne T, et al. In Vivo assessment of drug efficacy against Mycobacterium abscessus using the embryonic zebrafish test system. Antimicrob. Agents Chemother. 2014;58(7):4054–4063.

77. Bernut A, Nguyen-Chi M, Halloum I, et al. Mycobacterium abscessus-Induced Granuloma Formation Is Strictly Dependent on TNF Signaling and Neutrophil Trafficking. PLoS Pathog. 2016;12(11):.

78. Deng Q, Harvie EA, Huttenlocher A. Distinct signalling mechanisms mediate neutrophil attraction to bacterial infection and tissue injury. Cell. Microbiol. 2012;14(4):517–528.

79. Ellett F, Elks PM, Robertson AL, Ogryzko N V., Renshaw SA. Defining the phenotype of neutrophils following reverse migration in zebrafish. J. Leukoc. Biol. 2015;98(6):975–981.

80. Colom B, Bodkin J V., Beyrau M, et al. Leukotriene B4-Neutrophil Elastase Axis Drives Neutrophil Reverse Transendothelial Cell Migration InVivo. Immunity. 2015;42(6):.

