## Supplemental data for "Roscovitine exacerbates *Mycobacterium abscessus* infection by reducing NADPH oxidase-dependent neutrophil trafficking"

**Supplemental methodology**

**Mycobacterial strains and Growth conditions**

Mycobacterium abscessus sensu stricto (Mabs strains CIP104536^T^ morphotypes smooth (S) and rough (R)), carrying pTEC27-tdTomato^1^, pTEC25-Wasabi^1^ or the bacterial luciferase gene cassette *luxCDABE*^2^*,* grown in liquid medium as described^1,2^. Mabs inoculates were prepared for all infection challenges following established protocols^3^. Inocula were determined by microinjection onto VCAT (Vancomycin, Colistin sulfate, Amphotericin B, and Trimethoprim) chocolate agar plates (BioMérieux, France).

**Zebrafish Husbandry**

Maintenance of adult mutant fish was approved under home office license (P1A4A7A5E). Zebrafish experiments were performed on *i)* larvae <5 days post-fertilization (dpf) or ii) larvae >5 dpf to standards set approved by the Direction Sanitaire et Vétérinaire de l’Hérault et Comité d’Ethique pour l’Expérimentation Animale de la région Languedoc Roussillon (reference CEEA-LR-1145). Experimental procedures were performed using the WT pigment-less nacre^4^ and the recently validated knockout *cftr*^sh540^ mutant^5^, along with the transgenic line BAC(mpx:eGFP)i114^6^ labelling neutrophils with GFP. For neutrophil reverse migration assay, the *Tg(mpx:gal4)sh267;Tg(UAS:kaede)i222* line^7,8^, a neutrophil-specific transgenic line expressing a photoconvertable pigment, which allows photoconverted neutrophils to be tracked was used. The number of animals used for each procedure was guided by pilot experiments or by past results^1,5,9,10^.

For zebrafish anaesthesia procedures, larvae are immersed in a 0.168 mg/mL Tricaine solution in E3 media. When required, larvae were cryo-anesthetized by incubation on ice for 10 min and then euthanized using an overdose of Tricaine (500 mg/L).

**Morpholino injection**

Morpholino were purchased from Gene Tools. The splice-blocking morpholino for *cftr* knock-down (5’-GACACATTTTGGACACTCACACCAA-3’) was prepared and injected into one-cell-stage zebrafish as previously described^9^. Morpholino for *duox2* knock-down (5’-AGTGAATTAGAGAAATGCACCTTTT-3’) was prepared and injected as described earlier^11^. A standard control morpholino (5'-CCTCTTACCTCAGTTACAATTTATA-3') was used as a negative control.

**Inflammation assays in zebrafish larvae**

Inflammatory response was elicited by tail fin amputation (sterile inflammation) or local infection (infectious inflammation) on 3 dpf TgBAC(mpx:eGFP)i114 line according to procedures described earlier^5,6,9^. For sterile injury-induced inflammation assay, larvae were anesthetized then transection of the tail was performed with a microscalpel (5 mm depth; World Precision Instruments). For infection-induced inflammation assay, Mabs were locally injected into the muscle compartment of anesthetized larvae. Infected or injured larvae were maintained at 28°C in sterile E3 media until observations and analyzes by microscopy. Neutrophil chemotaxis was evaluated by assessing the number of cells at wound or infection sites at various time points throughout inflammatory processes on a fluorescence dissecting stereomicroscope (Leica). Larvae were mounted in 0.8% agarose supplemented with Tricaine then neutrophils at the area of interest were imaged on an Eclipse TE2000 U inverted compound fluorescence microscope (Nikon UK Ltd., Kingston upon Thames, UK) with a 10x or 20x NA objective lens.

**Pharmacological treatments for anti-inflammation and pro-resolution assays**

Larvae were incubated in sterile E3 media supplemented with 30 μM of R-Roscovitine (Roscovitine, gift from L Meijer, ManRos Therapeutics), 25 μM Tanshinone IIA (TIIA, Sigma-Aldrich)^12^, 100 μM Diphenyleneiodonium (DPI, Sigma-Aldrich)^5^, 1% Dimethyl sulfoxide (DMSO, Sigma-Aldrich) as negative controls. For anti-inflammation assays, prior to amputation/infection procedures, larvae were pre-treated with indicated compounds, then injured/infected and immediately put back in treatments. Neutrophils at wounds were counted at 4 hours post amputation (hpA) or 4 hours post infection (hpi) (neutrophil recruitment peaking) as previously observed^5,13^. For resolution of inflammation procedures, injured larvae were raised to 4 hpA then exposed with indicated drugs. Neutrophils at the wound sites were enumerated at 8 hpA for inflammation resolution as previously described^5^.

**In vivo Oxidative activity assay**

Epithelial oxidative response post-infection or injury was detected using CellROX® deep red (ThermoFisher) following established protocols^5,9^. Briefly, living zebrafish larvae were soaked in CellROX® reagent prior tail fin amputation procedure, then immediately injured and prepared for microscopy observations and analyses. ROS production at the wound sites was imaged by confocal microscopy (PerkinElmer Life) at 30 minutes post-amputation (mpA), and assessed in ImageJ 1.8 (National Institutes of Health) using the intensity of fluorescence and normalized to uninjured animal (WT or CF fish).

***In vivo* neutrophil reverse migration assay**

Sterile-inflammation was elicited by tail fin transection on 3 days post-fertilization (dpf) *Tg(mpx:gal4)sh267;Tg(UAS:kaede)i222* larvae following established methods^5^. Briefly, injured-larvae were raised to 4 hpA, then neutrophils at the wound sites was photoconverted by 120 pulses of the 405-nm laser at 40% laser power using an UltraVIEW PhotoKinesis device on an UltraVIEW VoX spinning disk confocal microscope (PerkinElmer Life and Analytical Sciences). Larvae were transferred to an Eclipse TE2000-U inverted compound fluorescence microscope (Nikon UK Ltd., Kingston upon Thames, UK), then were time-lapsed with 2.5 min intervals between 5 and 9 hpA using a 1394 ORCA-ERA camera (Hamamatsu Photonics Inc). Tracking analysis of red-fluorescing neutrophils moving away from photoconverted site was assessed in Volocity 6 (Improvision; PerkinElmer Life and Analytical Sciences), using the intensity of fluorescence to identify individually labeled neutrophils over the time course of inflammation resolution.

**Neutrophil apoptosis assays**

Rates of apoptotic neutrophils at wounds were assessed in 4% paraformaldehyde-fixed larvae after dual staining with Rhodamine-TUNEL (ApopTag Red; Millipore Corp.) to label apoptotic cells and FITC-TSA (TSAplus kit; Fluorescence Systems, PerkinElmer Life and Analytical Sciences) to label neutrophils as previously described^7^. Percentage of neutrophil apoptosis was performed at 8 hpA by confocal microscopy (PerkinElmer Life) and calculated by comparing the number of apoptotic neutrophils (dual TSA/TUNEL-positive) at the wound versus total mobilized neutrophils (TSA-positive).

***In vivo* tissue repair assay**

Regeneration assay was performed following established methods^5^. Briefly, 2 dpf embryos were Tricaine-anesthetized then tail fin transected. Tissue repair performances were evaluated by assessing the regenerated tail fin area at 3 dpA on an Eclipse TE2000 U inverted compound fluorescence microscope (Nikon UK Ltd., Kingston upon Thames, UK) with a 10x NA objective lens. Percentage of regeneration was calculated by normalizing the regenerated tail fin areas *versus* fin areas of unamputated animals (WT or CF fish).

**Systemic infection and treatment in zebrafish embryos**

Mabs were microinjected in zebrafish embryos as described earlier^14^. Briefly, systemic infections were carried out by the injection of single-cell suspensions of known titer (≈100-150 colony forming units (CFU), 2 nl) into the caudal vein of 30 hours post-fertilisation fish (hpf). Survival post-infection was assessed daily by counting dead embryos (no heartbeat) up to 10 days. To quantify fluorescent pixel count (FPC), which reflects the bacterial burdens, cords, abscesses, infected larvae were tricaine-anesthetized, mounted for real-time microscopy observations and processed as described^3^. To evaluated Roscovitine activity, larvae are incubated with Roscovitine at 1dpi. Water supplemented with Roscovitine was changed daily up to 4 days as previously described^15^.

**Mice infection and treatment**

The immunocompetent C3HeB/FeJ mouse model of Mabs infection^16^ was used to explore the therapeutic efficacy of Roscovitine. Briefly, mice were infected into the lateral caudal vein with 10^7^ CFU of R Mabs in 200 µl of water supplemented with 0.9% sodium chloride^17^. From 1 dpi, Roscovitine treatment was achieved by daily oral gavage^17^ (10 mg/kg or 50 mg/kg Roscovitine, respectively to 0.4 and 2 mg per mouse; 200 µl). A control group received an equivalent volume of 50% DMSO in polyethylene glycol 3000 solution (Sigma-Aldrich). To determine bacterial burden, mice were euthanized by cervical dislocation and lungs, spleen and liver from Roscovitine-treated and control mice were collected and homogenized, as previously described^18,19^. 10-fold serial dilutions were plated on VCAT chocolate agar plates and incubated for 5 to 6 days at 37°C prior to colony forming units (CFU) counts. The number of animals used was guided by pilot experiments or by past results^17^.

**Flow cytometry**

Mice were sacrificed and perfused through the left cardiac ventricle with PBS. Lungs were isolated, cut in small pieces and digested for 20 min at 37°C with 2.5 mg/ml Collagenase D (Roche). Cell suspensions were prepared and washed with complete RPMI medium (Gibco). Anti-CD11b (M1/70), anti-Ia-e (M5/114.15.2), anti-Ly6G (1A8) and anti-Ly6C (HK1.4) antibodies were purchased from Biolegend. Cells were stained with Fixable Viability Dye eFluor™ 780 (Thermofisher) then stained with antibody mix for 30 minutes at 4°C. Labeled cells were acquired on Fortessa flow cytometer (BD Biosciences) and data were analyzed using FlowJo software (TreeStar).

**Human cell infection and treatment**

Peripheral blood mononuclear cells (PBMC) were isolated from blood obtained from consented healthy volunteers and CF patients (approved by Regional NHS Research Ethics Committee) using Lympholyte®-H separation media (Cedarlane®, USA). Monocytes were extracted by CD14+ positive selection using magnetic beads (Miltenyi Biotec, UK), and the number of viable cells calculated using the trypan blue exclusion method. Macrophages were differentiated for 6 days in DMEM, 10% FCS, 100 U/mL penicillin/streptomycin, 2mM L-glutamine supplemented with 400 ng/mL M-CSF (Peprotech). Non-adherent cells were discarded prior to experiments.

Macrophage from healthy donors or CF patients were infected with bioluminescent Mabs then incubated with 25 µM Roscovitine or DMSO for 24 hours. At each specified time-point, cells were, washed and lysed then viable intracellular bacteria quantified as relative luminescent units (RLU) using a Glomax® 96 Microplate Luminometer.

**Lysosomal acidification activity**

THP1 cells were treated with DMSO or Roscovitine (25 μM) for 24 hours, washed and incubated for 15 minutes with LysoTracker® (40 nM) and LysoSensorTM Yellow/Blue DND-160 (1 μM) (Life TechnologiesTM) in Opti-MEMTM (Thermo Fisher Scientific) at 37°C with 5% CO2. LysoTracker® and LysoSensorTM Yellow/Blue DND-160 were used to quantify lysosomes in a cell and measure the pH of acidic organelles reciprocally. Cells were washed and Leibovitz’s L-15 medium (Thermo Fisher Scientific) added for imaging. Images were obtained using a spinning disk confocal microscope (Andor Technology) and analysis was done using Imaris Image Analysis Software.

To evaluated lysosomal fusion with intracellular Mabs, THP1 were infected then treated with Roscovitine. Following 24 hours of treatment, cells were washed with PBS, fixed using 4% formaldehyde, and permeabilized with 0.1% Triton X. A rabbit polyclonal anti-V-ATPase A1 antibody (H-140) (Santa Cruz Biotechnology, UK) was used to visualize the lysosomal V-ATPase. Samples were mounted for microscopy using ProLong® Gold antifade reagent with DAPI.

**Minimum inhibitory concentrations assay**

Roscovitine susceptibility testing for Mabs was determined using the microdilution method, in cation-adjusted Mueller-Hinton broth (Difco), according to the Clinical and Laboratory Standards Institute (CLSI) guidelines. In addition, the susceptibility profile was also determined on LB (Luria-Bertani) agar plates (Invitrogen) supplemented with increasing concentrations of Roscovitine, as reported^15^. Serial 10-fold dilutions of Mabs actively growing culture were inoculated or plated then incubated at 37°C for 3 to 4 days. The MIC was defined as the minimum concentration required to inhibit 99% of the growth.

**Data Sharing Statement**

For original data, please contact

**References**

1.	Bernut A, Herrmann JL, Kissa K, et al. Mycobacterium abscessus cording prevents phagocytosis and promotes abscess formation. *Proc. Natl. Acad. Sci. U. S. A.* 2014;111(10):.

2.	Chang M, Anttonen KP, Cirillo SLG, Francis KP, Cirillo JD. Real-time bioluminescence imaging of mixed mycobacterial infections. *PLoS One*. 2014;9(9):.

3.	Bernut A, Dupont C, Sahuquet A, et al. Deciphering and imaging pathogenesis and cording of Mycobacterium abscessus in zebrafish embryos. *J. Vis. Exp.* 2015;2015(103):.

4.	Lister JA, Robertson CP, Lepage T, Johnson SL, Raible DW. Nacre Encodes a Zebrafish Microphthalmia-Related Protein That Regulates Neural-Crest-Derived Pigment Cell Fate. *Development*. 1999;126(17):3757–3767.

5.	Bernut A, Loynes CA, Floto RA, Renshaw SA. Deletion of cftr Leads to an Excessive Neutrophilic Response and Defective Tissue Repair in a Zebrafish Model of Sterile Inflammation. *Front. Immunol.* 2020;11:.

6.	Renshaw SA, Loynes CA, Trushell DMI, et al. Atransgenic zebrafish model of neutrophilic inflammation. *Blood*. 2006;108(13):3976–3978.

7.	Elks PM, Van Eeden FJ, Dixon G, et al. Activation of hypoxia-inducible factor-1α (hif-1α) delays inflammation resolution by reducing neutrophil apoptosis and reverse migration in a zebrafish inflammation model. *Blood*. 2011;118(3):712–722.

8.	Holmes GR, Anderson SR, Dixon G, et al. Repelled from the wound, or randomly dispersed? Reverse migration behaviour of neutrophils characterized by dynamic modelling. *J. R. Soc. Interface*. 2012;9(77):3229–3239.

9.	Bernut A, Dupont C, Ogryzko N V., et al. CFTR Protects against Mycobacterium abscessus Infection by Fine-Tuning Host Oxidative Defenses. *Cell Rep.* 2019;26(7):1828-1840.e4.

10.	Bernut A, Nguyen-Chi M, Halloum I, et al. Mycobacterium abscessus-Induced Granuloma Formation Is Strictly Dependent on TNF Signaling and Neutrophil Trafficking. *PLoS Pathog.* 2016;12(11):.

11.	Niethammer P, Grabher C, Look AT, Mitchison TJ. A tissue-scale gradient of hydrogen peroxide mediates rapid wound detection in zebrafish. *Nature*. 2009;459(7249):996–999.

12.	Robertson AL, Holmes GR, Bojarczuk AN, et al. A zebrafish compound screen reveals modulation of neutrophil reverse migration as an anti-inflammatory mechanism. *Sci. Transl. Med.* 2014;6(225):.

13.	Bernut A, Nguyen-Chi M, Halloum I, et al. Mycobacterium abscessus-Induced Granuloma Formation Is Strictly Dependent on TNF Signaling and Neutrophil Trafficking. *PLoS Pathog.* 2016;12(11):.

14.	Székely R, Cole ST. Mechanistic insight into mycobacterial MmpL protein function. *Mol. Microbiol.* 2016;99(5):831–834.

15.	Bernut A, Le Moigne V, Lesne T, et al. In Vivo assessment of drug efficacy against Mycobacterium abscessus using the embryonic zebrafish test system. *Antimicrob. Agents Chemother.* 2014;58(7):4054–4063.

16.	Bernut A, Herrmann JL, Ordway D, Kremer L. The diverse cellular and animal models to decipher the physiopathological traits of Mycobacterium abscessus infection. *Front. Cell. Infect. Microbiol.* 2017;7(APR):

17.	Le Moigne V, Raynaud C, Moreau F, et al. Efficacy of Bedaquiline, alone or in combination with imipenem, against Mycobacterium abscessus in C3HeB/FeJ mice. *Antimicrob. Agents Chemother.* 2020;64(6):.

18.	Catherinot E, Clarissou J, Etienne G, et al. Hypervirulence of a rough variant of the Mycobacterium abscessus type strain. *Infect. Immun.* 2007;75(2):1055–1058.

19.	Rottman M, Catherinot E, Hochedez P, et al. Importance of T cells, gamma interferon, and tumor necrosis factor in immune control of the rapid grower Mycobacterium abscessus in C57BL/6 mice. *Infect. Immun.* 2007;75(12):.

**Table 1. MICs of Roscovitine against *M. abscessus* using the microdilution method in cation-adjusted Mueller-Hinton broth or on LB agar plates.**

| Mabs | MH broth  (µM) | LB agar  (µM) |
| --- | --- | --- |
| R variant | >256 | 128-256 |
| S variant | >256 | 128-256 |

**Supplemental figures**

**Figure S1.** **Comparative analysis of the *in vivo* efficacy of Roscovitine and Tanshinone IIA *in vivo***

(A) Larvae were injured, treated with Roscovitine or TIIA from 4 hpA then stained with TUNEL/TSA. Neutrophil apoptosis quantification at 8 hpA (n= 15, Fisher *t*-test). (B) *cftr* MO *Tg(mpx:gal4)sh267;Tg(UASkaede)i222 TgBAC(mpx:EGFP)i114* larvae were injured and treated from 4 hpA with Roscovitine or TIIA. At 4 hpA, neutrophils at site of injury were photoconverted. Plot showing the number of photoconverted neutrophils leaving the wound over 4 hpc. Line of best fit shown is calculated by linear regression. *P-*value shown is for the difference between the 2 slopes (*n*= 12, performed as 3 independent experiments).

**Figure S2. Evaluation of the *in vivo* efficacy of Roscovitine for tissue repair**

(A-B) Larvae were injured and treated from 4 hpA with Roscovitine or TIIA. Regenerative performance after treatments. (A) Regenerated fin areas are measured at 3 dpA (*n*= 21, One-Way ANOVA with Tukey’s multiples comparison test). (B) Representative imaging of injured tail fin at 3 dpA (Scale bars, 200 μm).

**Figure S3. Roscovitine enhances the lysosomal acidic function of human macrophage infected with Mabs**

(A-B) THP1 cells were treated with Roscovitine. After 24 hours of treatment, the effect of Roscovitine on lysosomal numbers and lysosomal acidification were evaluated in uninfected cells using LysoTracker® and LysoSensorTM (Scale bar, 10 μm). Roscovitine-treated cells showed an increase in the number of lysosomes per cell from 19.08 ± 3.27 in DMSO-treated cells to 33.18 ± 3.79 in Roscovitine-treated cell. An increase in the percentage of acidified lysosomes was also found in Roscovitine-treated cells (61.3 ± 4.68) compared to the DMSO control (41.97% ± 4.87). This graph represents the mean ± SEM of three independent experiments performed in triplicate. (C-D) THP1 cells were infected with Mabs expressing *wasabi* at a MOI 10:1 for 2 hours then treated with Roscovitine or DMSO. Following 24 hours of treatment, the effect of Roscovitine on lysosomal recruitment to Mabs was evaluated by analyzing co-localization of the lysosomal ATP dependent proton pump V-ATPase with intracellular Mabs. (D) Enhanced co-localization of lysosomal V-ATPase with Mabs was found in cells treated with Roscovitine compared to the DSMO control. Results represent the mean ± SEM of three independent experiments done in triplicate. Statistical significance: Fisher’s exact test of a contingency table (B) or two-tailed unpaired Student’s t test (D).

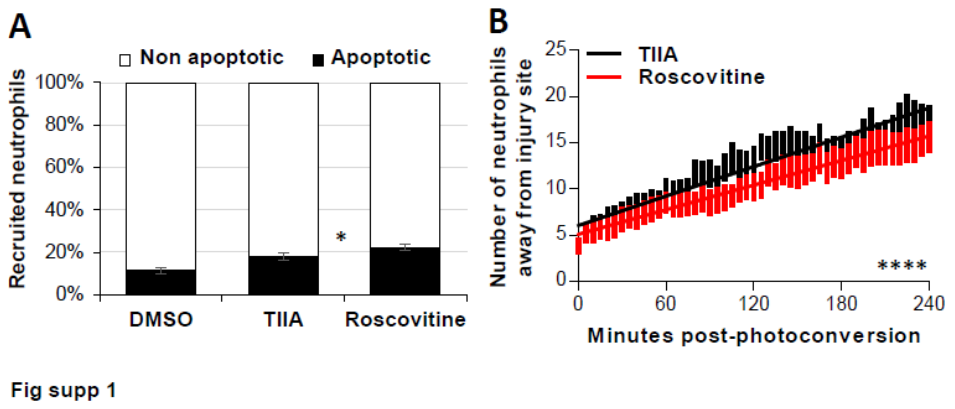

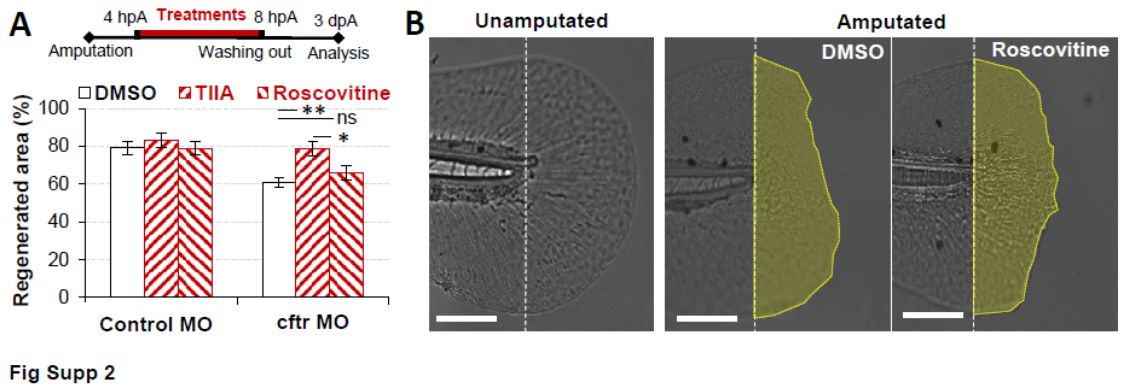

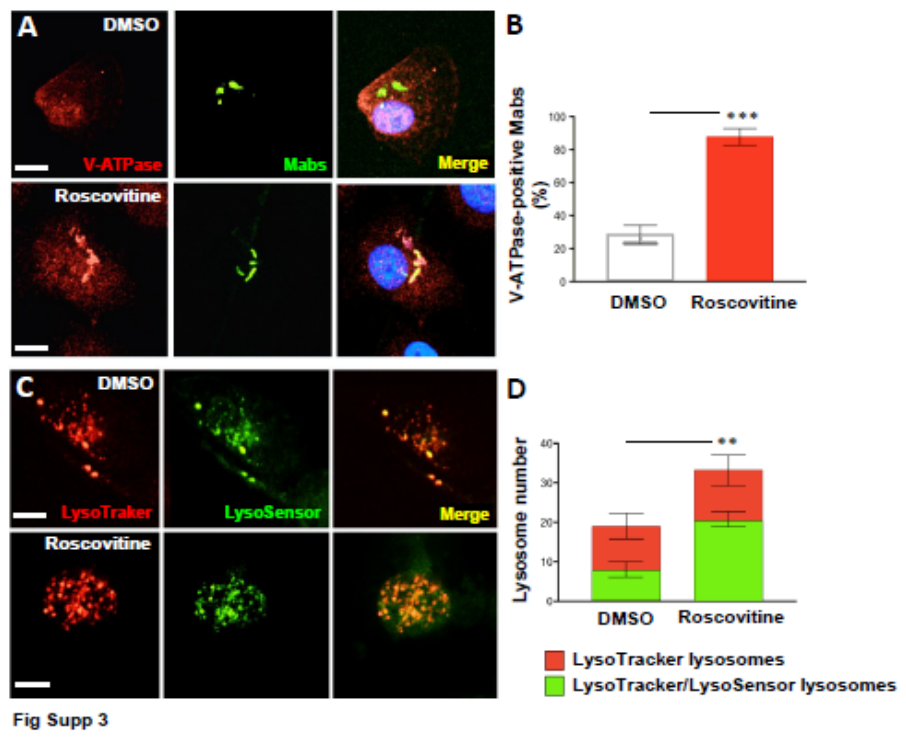
